# Visual gene expression reveals a cone to rod developmental progression in deep-sea fishes

**DOI:** 10.1101/2020.05.25.114991

**Authors:** Nik Lupše, Fabio Cortesi, Marko Freese, Lasse Marohn, Jan-Dag Pohlman, Klaus Wysujack, Reinhold Hanel, Zuzana Musilova

## Abstract

Vertebrates use cone cells in the retina for colour vision and rod cells to see in dim light. Many deep-sea fishes have adapted to their environment to have only rod cells in the retina, while both rod and cone genes are still preserved in their genomes. As deep-sea fish larvae start their lives in the shallow, and only later submerge to the depth, they have to cope with diverse environmental conditions during ontogeny. Using a comparative transcriptomic approach in 20 deep-sea fish species from eight teleost orders, we report on a developmental cone-to-rod switch. While adults mostly rely on rod opsin (*RH1*) for vision in dim light, larvae almost exclusively express middle-wavelength-sensitive (“green”) cone opsins (*RH2*) in their retinas. The phototransduction cascade genes follow a similar ontogenetic pattern of cone-followed by rod-specific gene expression in most species, except for the pearleye and sabretooth (Aulopiformes), in which the cone cascade remains dominant throughout development. By inspecting the whole genomes of five deep-sea species (four of them sequenced within this study: *Idiacanthus fasciola, Chauliodus sloani*; Stomiiformes; *Coccorella atlantica,* and *Scopelarchus michaelsarsi*; Aulopiformes), we found that deep-sea fish possess one or two copies of the rod *RH1* opsin gene, and up to seven copies of the cone *RH2* opsin genes in their genomes, while other cone opsin classes have been mostly lost. Our findings hence provide molecular evidence for a limited opsin gene repertoire and a conserved vertebrate pattern whereby cone photoreceptors develop first and rod photoreceptors are added only at later developmental stages.

## INTRODUCTION

Vision is a primary sense used by most vertebrates for navigation, predator avoidance, communication and to find food and shelter. At its initiation, vertebrate vision is enabled by cone (photopic, colour vision) and rod (scotopic) photoreceptors in the retina containing a light absorbing pigment that consists of an opsin protein covalently bound to a vitamin-A-derived chromophore (Lamb 2013). The absorbance of photons by the chromophore leads to a conformational change of the opsin protein, which initiates a photoreceptor-specific G-protein-coupled phototransduction cascade, propagating the signal to the brain (Downes and Gautam 1999, Larhammar et al. 2009, Lamb et al. 2019). It is thought that the development of the visual system follows a conserved molecular pattern whereby cone specific genes are activated first before the rod molecular pathway is initiated later during ontogeny (Mears et al. 2001, Shen and Raymond 2004, Sernagor et al. 2006). However, whether this is the case for all vertebrates and especially for those that have retinas that contain only rods as adults, remains unclear.

Changes in the light environment, ecology, and phylogenetic inertia are thought to be primary drivers for visual system diversity in vertebrates (Hunt et al. 2014). For example, most mesopelagic deep-sea fishes (200 – 1,000 m depth), either living strictly at depth or migrating to the shallows at night, have evolved visual systems that are sensitive to the dominant blue light (~ 470 – 490 nm) of their environment (Turner et al. 2009). Moreover, as the daylight and the bioluminescent light emitted by deep-sea critters are quickly dimmed with depth and distance, deep-sea fish visual systems have evolved peculiar morphologies to maximise photon capture including barrel-eyes, reflective tapeta and the use of rod-dominated and in many cases rod-only retinas that might be stacked into multiple banks (reviewed in de Busserolles et al. 2020). However, most mesopelagic fishes start their lives in the shallow well-lit epipelagic zone (0 – 200 m depth) (Moser and Smith 1993, Sassa and Hirota 2013). Consequently, their visual systems must cope with a variety of light intensities and spectra throughout development.

Studies investigating the gene expression in the retina of deep-sea fishes are scarce and usually focus on a selected few species (Zhang et al. 2000, Douglas et al. 2016, de Busserolles et al. 2017, Musilova et al. 2019a, Byun et al. 2020). In adults, species with pure rod retinas tend to only express rod opsin(s) (Douglas et al. 2016, Musilova et al. 2019a), albeit two species of pearlsides (*Maurolicus* spp.) have been found to express cone-specific genes (i.e., cone transduction pathway and opsin genes) inside rod-looking cells (de Busserolles et al. 2017). It remains unknown whether deep-sea fishes that have a low proportion of cone photoreceptors as adults (e.g., Munk 1990, Collin et al. 1998, Bozanno et al. 2007, Pointer et al. 2007, Biagioni et al. 2016) also express cone-specific genes at any stages of their lives or whether these fishes rely on the rod machinery alone. To investigate whether the retinal development in deep-sea fishes follows a similar cone-to-rod molecular pathway as found in other vertebrates or whether some species start their lives with the rod pathway activated, we set out to sequence the retinal transcriptomes of 20 deep-sea fish species, including the larval stages in ten species, belonging to eight different teleost orders (Argentiniformes, Aulopiformes, Beryciformes, Myctophiformes, Pempheriformes, Scombriformes, Stomiiformes and Trachichthyiformes). We have further investigated the genomic repertoire in five selected species.

## RESULTS AND DISCUSSION

### Opsin gene repertoire in the genome

In teleost fishes, gene duplications and deletions followed by functional diversification have resulted in extant species having between 1-40 visual opsin genes within their genomes (Musilova et al. 2019a, Musilova et al. 2021). These genes are defined by their photoreceptor specificity, their phylogeny, and their spectrum of maximal sensitivity (λ_max_) and fall within five major classes, four cone opsins (‘ultraviolet or UV sensitive’ *SWS1*: 347–383 nm, ‘blue’ *SWS2*: 397–482 nm, ‘green’ *RH2*: 452–537 nm and ‘red’ *LWS*: 501–573 nm) and one rod opsin (‘blue-green’ rhodopsin, *RH1* or Rho: 447–525 nm) (Carleton et al. 2020). We analysed the whole genomes of five deep-sea species (sawtail fish *Idiacanthus fasciola*, viperfish *Chauliodus sloani*; both Stomiiformes; sabretooth *Coccorella atlantica,* pearleye *Scopelarchus michaelsarsi*; both Aulopiformes; and fangtooth *Anoplogaster cornuta*; Trachichthyiformes), four of them sequenced for the purpose of this study. All species possess one or two copies of the rod opsin *RH1* gene, and one to seven copies of the *RH2* cone opsin (Fig. 1). All other cone opsin classes, i.e., the *SWS1*, *SWS2* (except for the fangtooth) and *LWS* are missing and have been putatively lost during evolution in these five species. This is in accordance with the observation that the *LWS* gene abundance decreases with the habitat depth (Musilova et al. 2019a). Such limited genomic repertoire most likely represents an evolutionary response to the deep-sea scotopic environment where the shortest (UV-violet) and longest (red) wavelengths of light get filtered out first in the water column, as opposed to middle-range wavelengths that can penetrate greater depths (reviewed in Musilova et al. 2021, De Busserolles et al. 2020, and Carleton et al. 2020). The increased *RH2* diversity observed in the two aulopiform species, on the other hand, illustrates the versatility of this cone opsin class and confirms its dominance in various dimmer-light habitats (Musilova and Cortesi 2021). Here we confirm that *RH2* is undoubtedly the most important (and often the only) cone opsin gene present in deep-sea fish genomes.

**Fig. 1:**
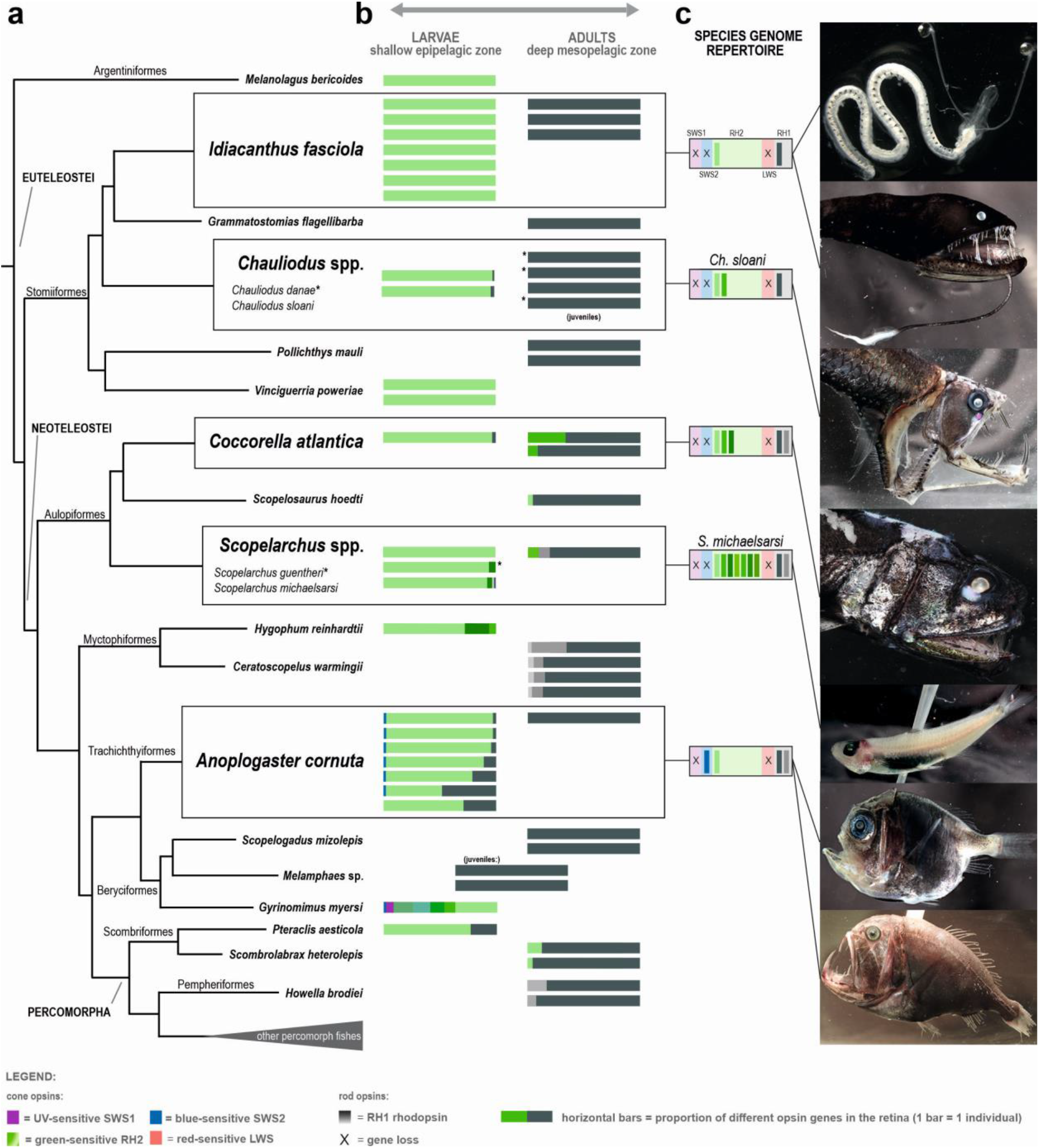
Cone and rod opsin gene expression in larval and adult deep-sea fishes. A) Transcriptomes were used to characterise the opsin gene expression in the retinas of 20 deep-sea fish species, belonging to eight different orders and mapped onto a simplified teleost phylogeny (topology after Betancur et al. 2017). Boxes highlight the five species for which both larval and adult samples were available. B) Proportional opsin gene expression for each individual (horizontal bar) at different developmental stages. Different colours correspond to cone (colours) or rod (shades of grey) opsin genes, depicted as the proportional expression over the total sum of visual opsins expressed. Different shades of the same colour represent multiple copies of the same gene family. Based on the opsin gene expression, the larvae (left column) show a pure-cone or cone-dominated retina, while the adults (right column) have a pure-rod or rod-dominated visual system. Juvenile specimens in two species had an adult expression profile. Note that some species expressed multiple RH1 copies (Scopelarchus, Howella brodiei and Ceratoscopelus warmingii adults) or multiple RH2 copies (Gyrinomimus sp. larva, Hygophum reinhardtii larva). Notably, adults and larvae of Scopelarchus sp. and Coccorella atlantica expressed different copies of RH2 (more details in Fig. 2). Details about the samples and expression levels are listed in Table 1. C) The genomic repertoire of the visual opsins is shown for five species Idiacanthus fasciola, Chauliodus sloani, Coccorella atlantica, Scopelarchus michaelsarsi (all this study), and Anoplogaster cornuta (Musilova et al. 2019a). The rod RH1 opsin and the cone RH2 opsin genes are present in all studied species in one or multiple (up to seven) copies. The SWS2 opsin gene was found only in the fangtooth, and the SWS1 and LWS are missing from all five studied genomes.

### Visual opsin gene expression

Transcriptomic sequencing of 20 deep-sea teleost species revealed that deep-sea fishes mainly express rod opsins and/or green-sensitive cone opsins (*RH2*s) in their retinas (Fig. 1, Table 1). While larvae mostly expressed *RH2*, adults and juveniles mostly expressed *RH1* and in a few cases a combination of both. We found none or very low expression of any of the other cone opsin genes: the red sensitive *LWS* was not expressed at all, the UV sensitive *SWS1* was only found in the larva of the whalefish, *Gyrinomimus* sp. (Beryciformes), and the blue/violet sensitive *SWS2* only in the larvae of the whalefish, and the fangtooth, *Anoplogaster cornuta* (Trachichthyiformes), (Fig. 1, Table 1). Differences in gene expression patterns are likely to be driven by ontogenetic transitions in light habitat from bright to dim environments and by changes in ecological demands, as discussed in more detail below.

**Table 1:**
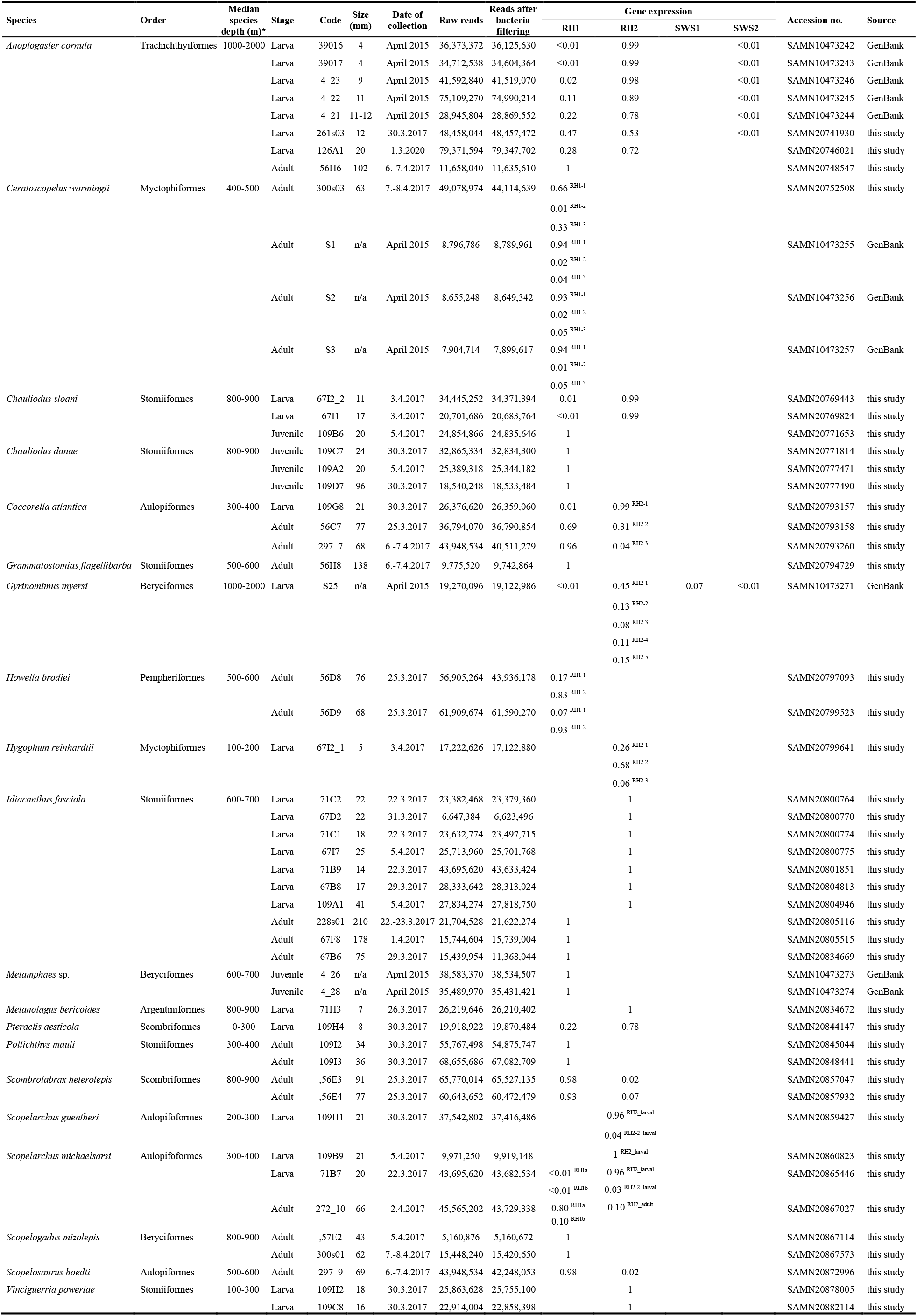
Samples used in the study and results of the opsin gene expression in the eyes or retina. * https://obis.org/

Similar to the opsin genes, we also detected ontogenetic differences in the expression of phototransduction cascade genes (Fig. 2). Here we focused on the comparison of five species from three teleost orders for which we had both larval and adult specimens available and found that the cone-specific genes were mostly expressed in the larval stages (e.g., cone transducin, *GNAT2*), while adults from three species mostly expressed rod-specific genes (e.g., rod transducin, *GNAT1*; Fig. 2b). Hence, at the molecular level, the visual systems of deep-sea fishes start out with a cone-based expression pattern. Moreover, in the fangtooth, where samples from various sized specimens were available, we found that the cone-specific expression was gradually replaced with the rod profile as the fish grew (Fig. 2c, Table S1). This sequence is similar to the visual development in shallower living fishes (e.g., Atlantic cod (Valen et al. 2016), zebrafish (Sernagor et al. 2006)) and terrestrial vertebrates (e.g., mice (Mears et al. 2001), rhesus monkey (La Vail et al. 1991)), where cone photoreceptors are first to develop, followed by temporally and spatially distinct rods (Raymond 1995, Shen and Raymond 2004). The cone-to-rod developmental sequence is therefore likely to be shared across vertebrates, even in species that have pure rod retinas as adults.

**Fig. 2:**
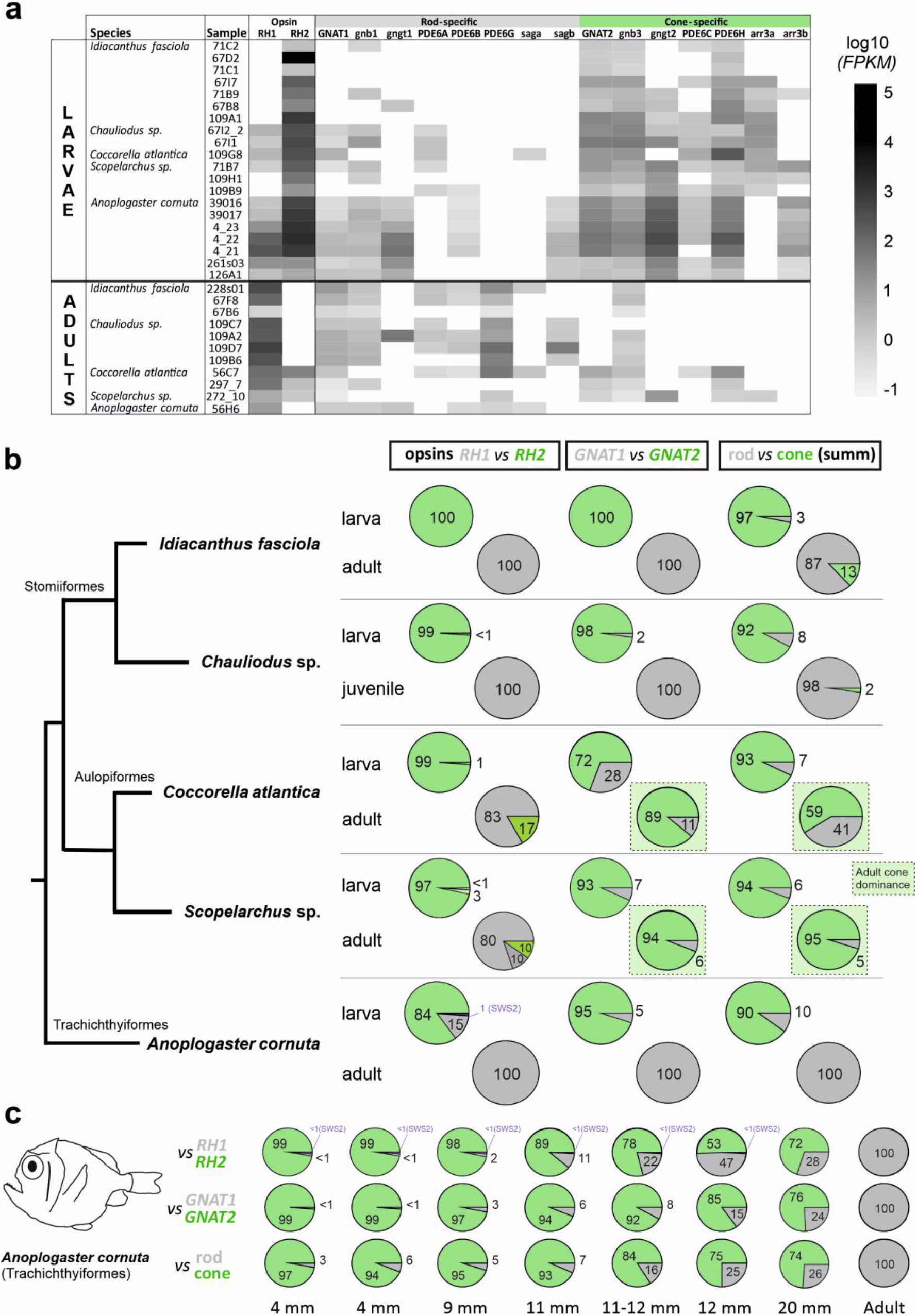
Phototransduction cascade gene expression in the retinas of five deep-sea fish species. A) Heat map of the expression of individual phototransduction cascade genes for each sample, based on normalised numbers of reads (FPKM). B) Pie charts comparing mean values of relative expression of the opsin genes (rod RH1 and cone RH2), photoreceptor-specific cascade transducin genes (rod-type GNAT1 and cone-type GNAT2), and all cascade genes (photoreceptor-specific transducins, arrestins and phosphodiesterases) summarized. The black square highlights the two aulopiform species with the discordance between the opsin type (rod-specific) and phototransduction cascade genes (cone-specific) in adults. C) Focus on the common fangtooth (Anoplogaster cornuta) transitional phase shown as a sequence for seven larval and one adult sample. Size given as standard length (SL). Note that all fangtooth larvae expressed both RH1 and RH2, with an increasing proportion of RH1 to RH2 as the larvae increased in size (with the exception of the largest larva where RH1:RH2 was 28:72). Smaller larvae also expressed the SWS2 gene. These individuals all had traits of larval phenotypes (dorsal and ventral horns and small teeth; Fig. 1) and were collected relatively shallow between 0–300 m using the plankton trawls.

### Ontogenetic shift in expression profiles and the transition phase

The observed developmental changes in the visual system are best explained by the different habitats larval and adult deep-sea fishes inhabit. In general, deep-sea fish larvae live in the shallow epipelagic zone (Moser and Smith 1993) where ambient light levels are sufficiently high to warrant a cone-based visual system. After metamorphosis, deep-sea fishes start to submerge deeper and take up a life at different depths in the mesopelagic or even bathypelagic (below 1,000 m depth) zone, where the sun- and moonlight is gradually replaced by bioluminescence as the main source of light (Denton 1990). In this extremely dim environment, rods work at their best and cone photoreceptors would be obsolete for the most part at least. Rod-based vision is also favoured in those deep-sea species that exhibit diel vertical migrations to feed in the plankton rich surface layers at night (de Busserolles et al. 2020). In addition, we discovered that in some species there was a switch in the expressed type of cone *RH2* opsin (Fig. 1). For example, in Aulopiformes, the larvae expressed an alternative *RH2* copy that is presumably sensitive to longer wavelengths of light compared to the *RH2* that was found in adults (Table 2). This clearly shows that larval and adult deep-sea fishes rely on different opsin expression profiles, which is similar to ontogenetic changes in opsin gene expression in diurnal shallow-water fishes such as freshwater cichlids (Carleton et al. 2016) and coral reef dottybacks (Cortesi et al. 2015, Cortesi et al. 2016) or between the freshwater and deep-sea maturation stages in eels (Zhang et al. 2000).

**Table 2:** Key-tuning amino acid sites in the cone opsin RH2 gene

| Species | Order | 83 | 122 | 207 | 255 | 292 | $\lambda_{\text{max}}$ (nm) | Reference |
| --- | --- | --- | --- | --- | --- | --- | --- | --- |
| Bovine RH1 |  | D | E | M | I | A | 500 | Yokoyama 2008 |
| Ancestral teleost |  | D | Q | M | I | A | 488 | Yokoyama & Yia 2020 |
| *Melanolagus bericoides* | Argentiniiformes | G | Q | . | V | . |  |  |
| *Coccorella atlantica* adult | Aulopiiformes | G | Q | . | . | . |  |  |
| *Coccorella atlantica* larval | Aulopiiformes | G | Q | . | V | . |  |  |
| *Scopelarchus michaelisarsis*_RH2_adult | Aulopiiformes | ? | Q | I | C | T |  |  |
| *Scopelarchus guentheri*_RH2_larval | Aulopiiformes | G | Q | . | V | . |  |  |
| *Scopelarchus michaelisarsis*_RH2_larval | Aulopiiformes | G | Q | . | V | . | 505 | Pointer et al. 2007 |
| *Scopelarchus guentheri*_RH2-2_larval | Aulopiiformes | G | Q | . | C | . |  |  |
| *Scopelarchus michaelisarsis*_RH2-2_larval | Aulopiiformes | G | Q | . | C | . |  |  |
| *Scopelosaurus hoedti* | Aulopiiformes | G | Q | . | . | . |  |  |
| *Gyrinomimus myersi* RH2-1 | Beryciiformes | G | Q | . | F | . |  |  |
| *Gyrinomimus myersi* RH2-2 | Beryciiformes | G | Q | L | F | . |  |  |
| *Gyrinomimus myersi* RH2-3 | Beryciiformes | G | Q | L | F | . |  |  |
| *Gyrinomimus myersi* RH2-4 | Beryciiformes | G | Q | L | F | . |  |  |
| *Gyrinomimus myersi* RH2-5 | Beryciiformes | G | Q | L | F | . |  |  |
| *Lepisosteus platyrhincus* (shallow outgroup) | Lepisosteiformes | G | . | . | . | . |  |  |
| *Hygophum reinhardtii* RH2-1 | Myctophiiformes | G | Q | . | V | . |  |  |
| *Hygophum reinhardtii* RH2-2 | Myctophiiformes | G | Q | . | V | . |  |  |
| *Hygophum reinhardtii* RH2-3 | Myctophiiformes | G | Q | . | V | . |  |  |
| *Lepidopus fitchi* RH2-1 | Scombriformes | G | . | . | V | . | 496 | Yokoyama & Yia 2020 |
| *Lepidopus fitchi* RH2-2 | Scombriformes | G | Q | . | V | . |  |  |
| *Lepidopus fitchi* RH2-3 | Scombriformes | G | Q | . | V | . | 506 | Yokoyama & Yia 2020 |
| *Lepidopus fitchi* RH2-4 | Scombriformes | G | Q | . | V | . |  |  |
| *Pteraclis aesticola* | Scombriformes | G | Q | L | . | . |  |  |
| *Scombrilabrax heterolepis* | Scombriformes | G | Q | . | . | . |  |  |
| *Aristostomias scintillans* | Stomiiformes | G | Q | L | V | . | 468 | Yokoyama & Yia 2020 |
| *Chauliodus* sp. | Stomiiformes | G | Q | . | F | . |  |  |
| *Grammatostomias flagellibarba* | Stomiiformes | G | Q | . | . | . |  |  |
| *Idiacanthus fasciola* | Stomiiformes | G | Q | . | V | . |  |  |
| *Vinciguerrria poweriae* | Stomiiformes | G | Q | . | F | . |  |  |
| *Anoplogaster cornuta* | Trachichthyiformes | G | Q | . | . | . |  |  |

Our data furthermore suggest that the ontogenetic change in visual gene expression precedes morphological changes such as metamorphosis from larva to juvenile and also habitat transitions. For example, in the fangtooth, the larvae which were collected from the shallows (0 – 300 m) showed increasing amounts of *RH1* expression with growth, despite displaying larval phenotypes throughout (horns and small teeth; Fig. 1). A similar pattern of changing opsin gene expression ahead of metamorphosis has also been reported from shallow-water fishes such as European eels (Bowmaker et al. 2008), dottybacks (Cortesi et al. 2016) and surgeonfishes (Tettamanti et al. 2019). Interestingly, all our fangtooth larvae (including the smallest individual with a total length of 4 mm) already expressed a small amount of *RH1* (Fig. 2c). Whether fangtooth start their lives with a pure cone retina or low-levels of rod opsin expression are normal even in pre-flexation larvae remains therefore unclear. In addition to the green-sensitive cone opsin *RH2*, the smallest fangtooth larvae also expressed low levels of the blue-sensitive *SWS2*, potentially conferring dichromatic colour vision to the early life stages of this species (Fig. 1).

### Photoreceptor cell identities

Interestingly, two aulopiform species; the Atlantic sabretooth, *Coccorella atlantica*, and the Bigfin pearleye, *Scopelarchus michaelsarsi*, despite expressing mostly *RH1* as adults, retained a cone-dominated phototransduction cascade expression profile akin to the one found in the larval stages (Fig. 2, Table S1). This begs the question whether the photoreceptors they are using are cones or rods in nature. Initially described in snakes and geckos (Simoes et al. 2016, Schott et al. 2019) and recently also in a deep-sea fish (de Busserolles et al. 2017), it appears that the dichotomy of rods and cones is not always as clear cut as one might think. For example, adult deep-sea pearlsides, *Maurolicus* spp. have a retina that expresses ~ 99% *RH2* and ~ 1% *RH1* with corresponding cone and rod phototransduction gene expressions. Their photoreceptors, however, are all rod-shaped and careful histological examination has shown that these consist of a tiny proportion of true rods and a majority of transmuted rod-like cones (de Busserolles et al. 2017). In the case of pearlsides, and also in geckos and snakes, the opsin and phototransduction genes correspond to each other making it possible to distinguish photoreceptor types at the molecular level. However, in the aulopiforms, high expression of rod opsin is seemingly mismatched with high levels of cone phototransduction gene expression (Fig. 2). In salamanders, the opposite pattern can be found whereby a cone opsin is combined with the rod phototransduction cascade inside a rod looking cone photoreceptor (Mariani 1986). Anatomically, the retina of *S. michaelsarsi* is composed of mostly rods with low numbers of cone cells (Collin et al. 1998), while the adult retina of *Evermanella balbo,* an evermannellid species related to *C. atlantica,* appears to consist of two differently looking rod populations (Wagner et al. 2019). It is therefore likely, as found in pearlsides (de Busserolles et al. 2017), that these fishes have a high proportion of transmuted rod-like cone photoreceptors, but that they use *RH1* instead of a cone opsin as the visual pigment. Alternatively, a proportion of true rods might make use of the cone phototransduction cascade. Either way, combining more stable rod opsin in a rod-shaped cell with the cone-specific cascade is likely to increase sensitivity while also maintaining high transduction and recovery speeds of cells (Baylor 1987, Kawamura and Tachibanaki 2012, Luo et al. 2020). Histology, fluorescent in-situ hybridisation and ideally physiological recordings are needed to ultimately disentangle the identity of photoreceptor cells in aulopiforms.

### Evolutionary history of deep-sea fish opsins

While the majority of adult fishes relied on a single *RH1* copy, we found three species that expressed multiple *RH1* copies: The Warming's lanternfish, *Ceratoscopelus warmingii* (Myctophiformes), expressed three different *RH1* genes, and *S. michaelsarsi* and the basslet, *Howella brodiei* (Pempheriformes), expressed two copies each. Larvae and a few adult deep-sea fishes mostly expressed a single *RH2* copy, except for the pearleyes, *Scopelarchus* spp., and the Reinhardt’s lanternfish, *Hygophum reinhardtii*, which expressed three larval copies each, and the whalefish (*Gyrinomimus* sp.) which expressed five larval copies (Fig. 1).

The *RH1* and *RH2* phylogenies revealed that most deep-sea fish visual opsins cluster together by species or order (Fig. 3). For example, in the whalefish all *RH2s* are clustered together suggesting that these genes are lineage or species-specific duplicates (Fig. 3b). However, there were a few exceptions, suggesting more ancient duplication events. In *Scopelarchus* the two *RH1* copies are not in a sister relationship and in fact result in different clusters, suggesting that these copies originated in the common ancestor of aulopiforms or perhaps even earlier (Fig. 3a). Also, the *RH2s* in aulopiforms (*Scopelarchus*, *Coccorella*) cluster by ontogenetic stage, making it likely that the developmental switch in gene expression was already present in the aulopiform ancestor (Fig. 3b).

**Fig. 3:**
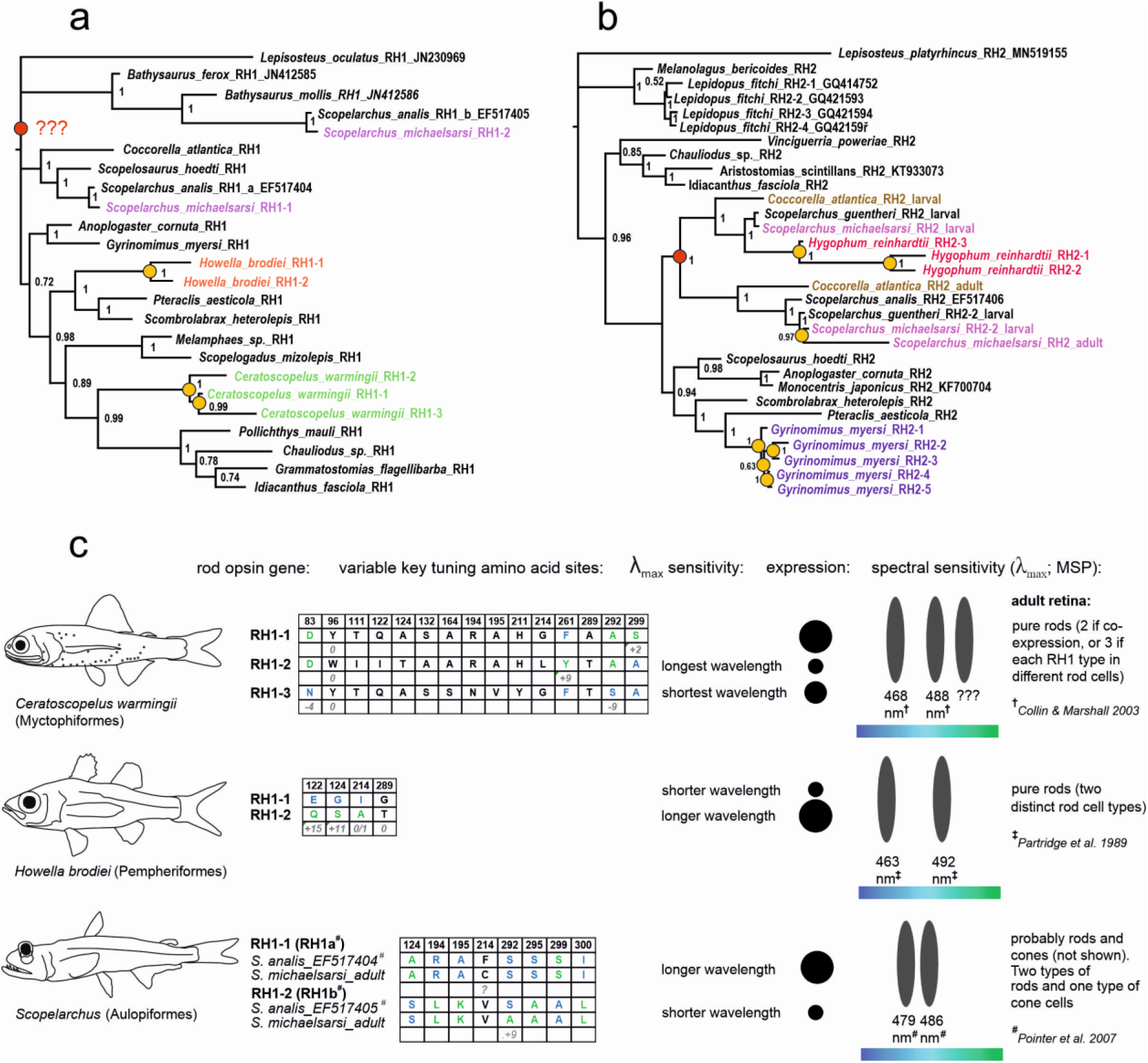
Gene trees of the A) RH1 and B) RH2 opsin genes found in the retinal transcriptomes of deep-sea fishes. Species with multiple copies are highlighted in colour. Additional gene sequences from public databases are listed with their GenBank accession numbers. Note the topology within Aulopiformes; the adult RH2s of Cocorrella atlantica and Scopelarchus cluster together as do the major larval RH2s. Yellow circles mark lineage-specific gene duplication events, while red circles pinpoint the ancestral duplication of RH1 impacting the Scopelarchus genus, and the duplication of RH2 in the aulopiform ancestor (or at least the common ancestor of Coccorella and Scopelarchus). C) Key tuning spectral site mutations in species with multiple rod opsins and indicative wavelength shifts based on the previous in vitro experiments. The known shifts are listed in nanometers. Blue and green letters in the tables stand for the shorter- and longer-shifting amino acid variants, respectively. Multiple different rod opsins have been found in three species, Ceratoscopelus warmingii (Myctophiformes), Howella brodiei (Pempheriformes) and Scopelarchus michaelsarsi (Aulopiformes). Note that the RH1 copies in Scopelarchus seem to show a mixed pattern - the longer-wavelength sensitive copy (RH1a; confirmed by in vitro measurements by Pointer et al. 2007) carries also several shorter-shifting amino-acid sites as compared to RH1b). For functional interpretation we considered microspectrophotometry measurements from # = Pointer et al. 2007, † = Collin and Marshall 2003 and ‡ = Partridge et al. 1989.

### Molecular complexity of deep-sea fish visual systems

The complexity of the deep-sea fish visual systems at the molecular level varied quite substantially. For example, the three Stomiiformes species: The Ribbon sawtail fish, *Idiacanthus fasciola*, and two species of viperfish, *Chauliodus sloani* and *Ch. danae*, appeared to have a very basic visual set up; these fishes were found to express a single *RH2* cone opsin as larvae and a single *RH1* rod opsin as adults (Fig. 1). On the contrary, several deep-sea fish orders examined here expressed more than one opsin gene. Adult lanternfishes and basslets have rod-only retinas but expressed multiple *RH1* copies that have functionally diversified (Fig. 3). Other species expressed both cone and rod opsins as adults (the aulopiform species and *Scombrolabrax*), which is somewhat similar to the opsin gene expression profiles found in shallow-living nocturnal reef fishes (Cortesi et al. 2020).

The most complex visual system in this study was found in *S. michaelsarsi*. In general, this species is known for its numerous morphological and anatomical adaptations to vision in the depth, including having barrel eyes with a main and an accessory retina, rods that are organised in bundles, large ganglion cells and corneal lens pads (Collin et al., 1998). The two copies of *RH1* (*RH1a* and *RH1b*) it expressed showed high sequence diversity differing in 79 out of 354 amino acids, eight of which are known key tuning sites likely to change the spectral sensitivity of the pigments via a shift in λ_max_ (Fig. 3, Table 3) (Yokoyama 2008, Musilova et al. 2019a, Yokoyama and Yia 2020). This supports the findings by Pointer et al. (2007) who found by using in vitro expression of pigments in another pearleye species, *S. analis*, two rod photoreceptors with different absorption maxima at 479 and 486 nm. Interestingly, Pointer et al. (2007) also speculate that another short-shifted opsin (previously measured in *Sc. analis* to have λ_max_ at 444 nm by Partridge et al., 1992) possibly belongs to the SWS2 class. Our data however does not support this expectation as no SWS2 gene is found in the genome of *Scopelarchus michaelsarsi*. The existence and identity of such short-sensitive opsins in pearleyes remains therefore elusive. The situation is less clear for the green-sensitive *RH2* opsin. While in *S. analis* cones have been found in the accessory and main retinas, in *S. michaelsarsi* cone photoreceptors appear restricted to the accessory retina alone (Collin et al. 1998). This is intriguing as it suggests substantial differences in visual systems even between closely related species from the same genus.

**Table 3:**
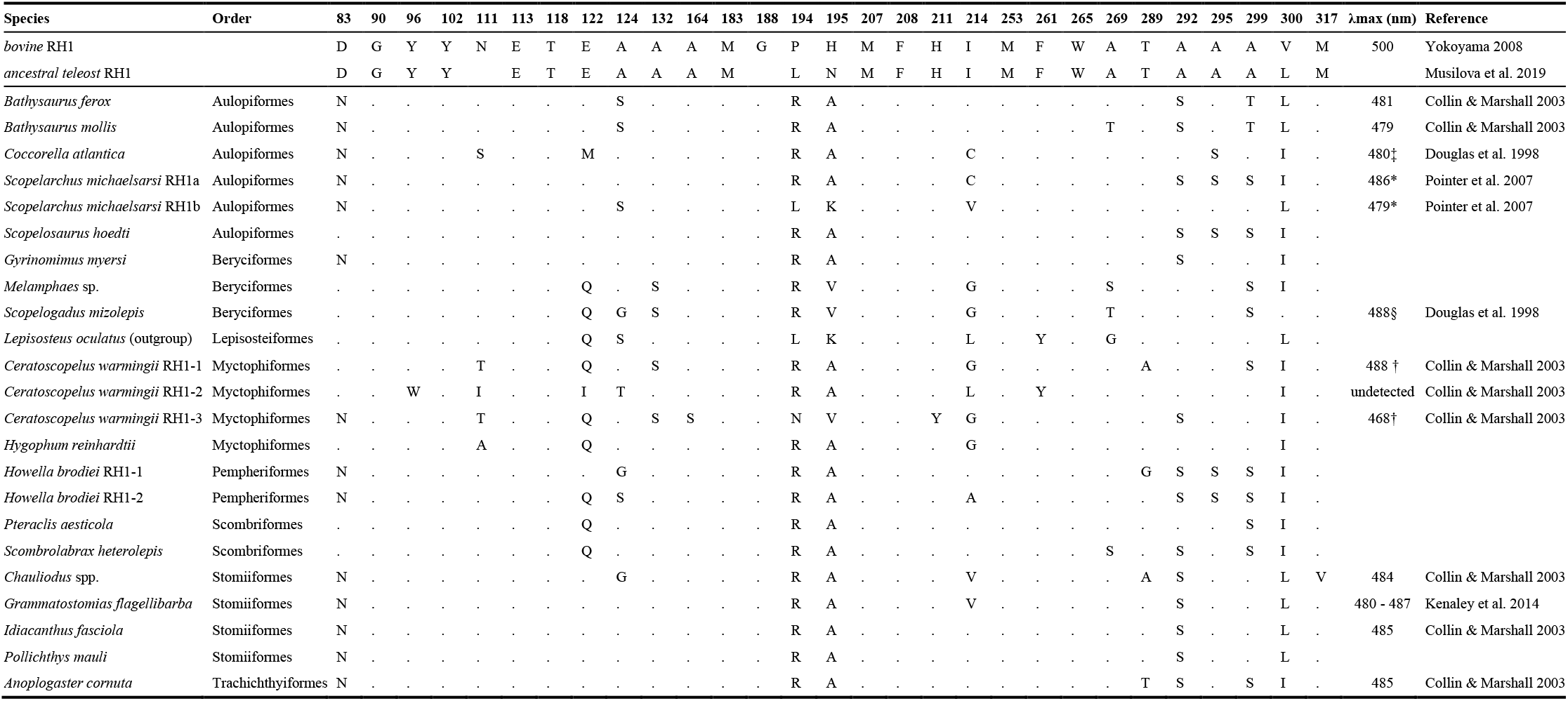
Key-tuning amino acid sites in the rhodopsin RH1 gene. * = value for Scopelarchus analis; † = two pigments reported without assignment to the gene; see also Figure 3; ‡ = value for closely related species, Evermannella balbo; § = value for Scopelogadus beani.

### The visual ecology of deep-sea fishes

We found molecular support for deep-sea visual adaptations on multiple levels:

#### 1) Opsin gene diversity in the genome

Data from the genomes of five deep-sea species revealed the diversity of the opsin genes (Fig. 1C). Retinal transcriptomes in the Stomiiformes pointed towards a simple visual system that is based on a single expressed opsin gene at different developmental stages (*RH2* in larvae, *RH1* in adults) (Fig. 1). A search for visual opsins in the stomiiform genomes sequenced for the purpose of this study (*I. fasciola* and *Ch. sloani*), as well as in several published genomes (Musilova et al., 2019a) revealed that Stomiiformes are likely to have lost all cone opsin gene families except for *RH2* (Fig. 1). In the case of *RH2*, they seem to only have a single or at most two gene copies, which is substantially less than other teleosts (Musilova et al. 2019a; Musilova and Cortesi, 2021). The stomiiform example, therefore, shows that a decrease in light intensity and the spectrum of light in the deep-sea may not only restrict gene expression at adult stages, but also lead to the loss of opsin genes altogether. Similarly, a loss in opsin and other vision-related genes (e.g., otx5b, crx) has previously been reported from shallow living fishes that are either nocturnal (Luehrmann et al. 2019), live in murky waters (Liu et al. 2019), or inhabit caves or similarly dim environments (Huang et al. 2019, Musilova et al. 2019a). Contrarily, the genomes of two aulopiform species (*C. atlantica* and *S. michaelsarsi*) revealed expanded cone opsin gene repertoires achieved mostly through *RH2* duplications (*C. atlantica* with three *RH2*, and *S. michaelsarsi* with seven *RH2* genes; Fig 1). Both species also possess two copies of the rod opsin gene. These species inhabit relatively shallower depths (300-400m) compared to other deep-sea fishes such as the Stomiiformes (Table 1). It is likely that having extra copies of *RH2* cone opsins may benefit their vision at these photon-richer depths. It is also possible that an evolutionary stochasticity and the gene content in the ancestor have contributed to the observed pattern. To be able to clearly state this, future research should be done on multiple aulopiform species.

#### 2) Visual gene expression

Previous work has found that the expression of the longest-(*LWS* - red) and shortest-(*SWS1* - UV) sensitive opsins is reduced or absent in deeper living coral reef fishes (Cortesi et al. 2020) and in fishes inhabiting deep freshwater lakes (e.g., Hunt et al. 1997, Sugawara et al. 2005, Musilova et al. 2019b), which is correlated with a loss of short- and long-wavelengths with depth. Also, many deep-sea fish lineages have lost *LWS* from their genomes (Musilova et al. 2019a, Cortesi et al. 2021). Supporting these findings, we show here that deep-sea fishes lack any *LWS* expression even in the shallow-living larval stages (Fig. 1). Similarly, *SWS1* is not expressed in any of the species studied, except for in the larval whalefish, and is also absent from many deep-sea fish genomes (Fig. 1) (Musilova et al. 2019a). However, shallow larval stages are likely to explain why all deep-sea fishes studied to date maintain at least some cone opsins in their genomes (Musilova et al. 2019a).

Most deep-sea fish larvae expressed a single *RH2* gene, but the larvae of some species (fangtooth, whalefish, and lanternfish) expressed multiple cone opsin genes (Fig. 1). This is likely to provide them with similar visual systems to the larvae of shallow-living marine (Britt et al. 2001) and freshwater species (Carleton et al. 2016), possibly aiding in detecting residual light and discriminating brightness and/or colours. Juvenile deep-sea fishes, on the other hand, showed rod-based expression profiles also found in the adult stages (Fig. 1). This shift in opsin gene expression correlates with developmental changes in ecology. As opposed to the adults which are exposed to a narrow and dim light environment where food is scarce, larvae typically live in well-lit, broad spectrum shallow waters where food and predators are abundant (Moser and Smith 1993).

#### 3) Functional adaptation in key spectral tuning sites

When multiple *RH1* copies were expressed, they often showed distinct differences in key amino acid sites that are likely to shift the spectral sensitivities of the pigments (Fig. 3 and Table 3) (Yokoyama 2008, Musilova et al. 2019a). Estimating spectral absorbance from these sites resulted in very similar λ_max_ values to those that were measured in vivo from the same or closely related species using microspectrophotometry (MSP) or similar techniques (Partridge et al. 1989, Collin and Marshall 2003, Pointer et al. 2007) (Fig. 3, Table 3). We therefore integrated the available functional evidence (MSP values) with our molecular data (protein sequence and gene expression levels) to better understand the function of the visual system in the three species that expressed multiple rod opsins (Fig. 3, Table 3).

The three different *RH1*s in *C. warmingii* differed in 15 key-tuning sites. Our data revealed a dominant rod opsin copy (*RH1-1*), and a shorter-(*RH1-2*) and a longer-shifted (*RH1-3*) copies with lower transcript levels (Figs. 1 and 3). The MSP in *C. warmingii* revealed two distinct rod types with λ_max_ values of 488 and 468 nm in past (Collin and Marshall 2003) corresponding most likely to the *RH1a/RH1-1* and *RH1b/RH1-3* genes, respectively. It is possible that multiple *RH1* copies are coexpressed within the same photoreceptor, something that has previously been reported for cone opsins in shallow-water marine (Savelli et al. 2018, Stieb et al. 2019) and freshwater fishes (Dalton et al. 2014, Torres-Dowdall et al. 2017). Coexpression could produce visual pigment mixtures that shift photoreceptor sensitivity and enhance visual contrast, aiding in predator-prey interactions or mate detection (Dalton et al. 2014). Alternatively, we predict that a third rod photoreceptor type with longer spectral sensitivity (*RH1-2*; Fig 3c) exists, possibly overlooked during MSP, which can happen especially if the cell type is rare. While the function of having multiple rod opsins in *C. warmingii* remains to be investigated, several possible benefits for a multi-rod visual system have recently been proposed including that it might enable conventional or unusual colour vision in dim light, it might be used to increase visual sensitivity, or enhance an object’s contrast against a certain background (Musilova et al. 2019a).

In *H. brodiei*, the second *RH1* copy (*RH1-2*) differed in two key-tuning sites, E122Q (−15 nm) and G124S (−11 nm), known to cause major short-wavelength shifts in other fishes (Yokoyama 2008). This is in accord with the MSP measurements in its sister species, *Howella sherborni*, which found two different rod types with spectral sensitivities of 463 and 492 nm (Fig. 3) (Partridge et al. 1989). Having multiple differently tuned rod photoreceptors, one centred on the prevailing light (bioluminescence and/or ambient light ~ 480 – 490 nm) and a second one that is offset from it (i.e., the offset pigment hypothesis; Lythgoe 1966), may be beneficial to break counter illumination of prey - a way of active camouflage in mesopelagic organisms where ventral photophores emit bioluminescent light that matches the residual down-welling light (Denton et all. 1985). Hence, revealing an individual’s silhouette could help to distinguish prey and predators from the background lighting, or visually finding mates. However, apart from lanternfishes with three (or more) and basslets with two rod opsins, or exceptional cases of tube-eye (six) and spinyfin (38), the majority of the deep-sea fishes seem to have only one rod opsin (Musilova et al., 2019a).

Differences in spectral sensitivity between photoreceptors can also be due to the chromophore type that binds to the opsin protein; visual pigments with a vitamin A1-based chromophore (typical in marine fishes) confer shorter-shifted λ_max_ values compared to those with a vitamin A2-based chromophore (typical in freshwater fishes) (Carleton et al. 2016). *Cyp27c1* is the enzyme responsible for converting A1- to A2-based chromophores (Enright et al. 2015), with high expression levels suggesting the presence of longer-shifted visual pigments. However, *Cyp27c1* was not expressed in our dataset suggesting that the visual pigments of these deep-sea fishes are based on A1-retinal alone (Table S2).

### Conclusions

So far, the development of deep-sea fish vision at the molecular level had not been studied in detail and only limited morphological information is available. In this study we compared opsin and visual gene expression between 20 deep-sea fish species revealing a major change in expression between larval and adult stages. While deep-sea fish genomes contain both cone and rod opsin genes, larvae rely on the cone pathway and most adult fishes switch to a rod-dominated or rod-only visual system. The cone- versus rod-specific phototransduction cascade genes follow the opsins in some lineages, however, not in aulopiforms. We detected reduced opsin gene repertoires in the genomes of five deep-sea fish species composed only of one rod (*RH1*) and one or two cone (*RH2*, or *RH2* and *SWS2*) opsin gene classes. Interestingly, we have discovered lineage-specific opsin gene duplications, possibly allowing for increased visual sensitivity and certain kind of colour vision in the depth in some species. Overall, our molecular results support a conserved developmental progression in vertebrates whereby cones appear first in the retina and rod photoreceptors are added later during development.

## MATERIALS AND METHODS

Specimens used in this study were collected in the Sargasso Sea during three multipurpose fishery surveys conducted by the German Thünen Institute of Fisheries Ecology onboard the research vessels *Maria S. Merian* in March to April in 2015, and *Walther Herwig III* in 2017 and in 2020. The sampling of adults occurred during both day and night at depths of 600 – 1'000 m using a mid-water pelagic trawl net (Engel Netze, Bremerhaven, Germany) with an opening of 30 m × 20 m, a length of 145 m, and mesh sizes (knot to knot) from 90 cm decreasing stepwise to 40, 20, 10, 5, 4, 3, 2 cm, with a 1.5-cm mesh in the 27-m-long codend. The larvae were mostly collected using an Isaacs-Kidd Midwater Trawl net (IKMT; 6.2 m^2^ mouth-opening, 0.5 mm mesh size; Hydro-Bios Apparatebau GmbH) at depths of 0 - 300 m by double-oblique transect tows. Adult fish were flash-frozen at −80 °C upon arrival on board and their fin clip was stored in 96% ethanol. Larval samples were fixed in RNAlater™ (ThermoFisher) and stored at −80 °C until further use.

To sequence the whole genome of *Idiacanthus fasciola, Chauliodus sloani, Coccorella atlantica,* and *Scopelarchus michaelsarsi,* the genomic DNA was extracted from the fin clip using the DNeasy Blood and Tissue kit (Qiagen) following the enclosed protocol. The library preparation and genome sequencing on Illumina NovaSeq platform (150 bp PE and the yield over 20 Gb per genome) has been outsourced to the sequencing centre Novogene, Singapore (https://en.novogene.com/). To analyse the opsin gene repertoire, the raw genomic reads were mapped in Geneious software version 11.0.3 (Kearse et al. 2012) against the opsin references (single exons of all five opsin classes from the reference species: Nile tilapia, Round goby, Blind cavefish, Spotted gar), as well as against the genes found in the transcriptomes of each species. The parameters were set to the Medium Sensitivity to capture all reads that matched any visual opsin gene. The captured reads mapping to all exons were then remapped against one reference per exon and the species-specific consensus sequence was generated. If present, multiple paralogous genes were disentangled manually, and the consensus sequence was exported for each variant (see below more details for the transcriptomic analysis). The obtained consensus sequences served as references for the second round of mapping, whereby all genomic reads were again mapped with the Low Sensitivity settings, and each reference was then elongated by the overlapping sequence. This step was repeated until the full gene region was covered. In case of *Scopelarchus michaelsarsi*, we were not able to cover the full length of five out of seven *RH2* genes due to the repetitions and these genes were reported in two parts, always one covering the exons 1 and 2, and one covering exons 3, 4 and 5. The genomic raw reads are available from GenBank (BioProject PRJNA754116) and the opsin gene sequences are provided in Supplementary file 1.

Total RNA was extracted from the whole eyes using either the RNeasy micro or mini kit (Qiagen) and the extracted RNA concentration and integrity were subsequently verified on a 2100 Bioanalyzer (Agilent). RNAseq libraries for 31 samples were constructed in-house from unfragmented total RNA using Illumina’s NEBNext Ultra II Directional RNA library preparation kit, NEBNext Multiplex Oligos and the NEBNext Poly(A) mRNA Magnetic Isolation Module (New England Biolabs). Multiplexed libraries were sequenced on the Illumina HiSeq 2500 platform as 150 bp paired-end (PE) reads. Library construction and sequencing (150 bp PE) for an additional 10 samples was outsourced to Novogene, Singapore (https://en.novogene.com/). We additionally re-analysed 11 retinal transcriptomes previously published in Musilova et al. (2019a). Together, then, our dataset comprised 53 samples of which, based on morphology, 26 were classified as larvae, 6 as juveniles and 21 as adults. Sample IDs, number of raw reads, individual accession numbers for BioProject PRJNA754116 and further parameters are listed in Table 1.

The sequence data was quality-checked using FastQC (Andrews 2010). Opsin gene expression was then quantified using Geneious software version 11.0.3 (Kearse et al. 2012). For each sample we first mapped the reads against a general fish reference dataset comprising all visual opsin genes from the Nile tilapia, *Oreochromis niloticus* and the zebrafish, *Danio rerio*, with the Medium-sensitivity settings in Geneious. This enabled us to identify cone and rod opsin specific reads. If present, paralogous genes were subsequently disentangled following the methods in Musilova et al. 2019a and de Busserolles et al. 2017. Briefly, we created species-specific references of the expressed opsin genes and their several copies (Musilova et al. 2019a) and re-mapped the transcriptome reads with Medium-Low sensitivity to obtain copy-specific expression levels. If multiple opsin genes were found to be expressed, we report their proportional expression in relation to the total opsin gene expression (Fig. 1). We used the same pipeline to quantify expression of phototransduction cascade genes in five focal deep-sea species (Fig. 2, Table S1), and to search for the expression of the *cyp27c1* gene (Table S2).

To analyse key amino-acid substitutions in *RH1* and *RH2* and potential shifts in their absorbance, we first translated the opsin coding sequences into amino acid sequences, and then aligned them with the bovine *RH1* (GenBank Acc.No: M12689). We have specifically focused on the positions identified as key-tuning sites in Yokoyama (2008) and Musilova et al. (2019a). For details, see Tables 2 and 3. Unfortunately, we were not able to estimate the sensitivity shift of rod opsin copies in *C. warmingii* as only four of the amino acids that were substituted at the 15 key-tuning amino acid sites corresponded with previously tested cases (Yokoyama, 2008, Musilova et al. 2019a). Out of the three copies, *RH1-2* has three out of four longer-shifting amino acid variants in these four sites and we assume it is therefore red-shifted. *RH1-1* is most likely sensitive to 488 nm, and *RH1-3*, being the shortest, to 468nm (Fig. 3c).

A dataset containing *RH1* opsin gene sequences mined from our dataset and additional *RH1*s obtained from GenBank (GenBank accession numbers listed in Figure 3), were aligned using the MAFFT (Katoh et al. 2009) plugin as implemented in Geneious, and a phylogenetic tree was subsequently reconstructed using MrBayes v3.2.1 (Ronquist and Huelsenbeck 2003) (Fig. 3a). Trees were produced using the Markov chain Monte Carlo analysis which ran for 1 million generations. Trees were sampled every 100 generations, and the printing frequency was 1000, discarding the first 25% of trees as burn-in. The evolutionary model chosen was GTR model with gamma-distributed rate variation across sites and a proportion of invariable sites. Posterior probabilities (PP) were calculated to evaluate statistical confidence at each node. We used the same approach with an *RH2*-specific reference dataset to reconstruct the phylogenetic relationship between the transcriptome-derived deep-sea *RH2* genes (Fig. 3b).

## Acknowledgements

We would like to express our thanks to both scientific and technical crew of the Maria S. Merian and Walther Herwig III research cruises in 2015, 2017 and 2020. In addition, we thank Tina Blancke for help with the sample management, and Veronika Truhlářová for technical support and lab management. We would also like to thank three anonymous reviewers for their comments improving the final version of the manuscript. NL and ZM were supported by the Swiss National Science Foundation (PROMYS - 166550), ZM by the PRIMUS Research Programme (Charles University), the Czech Science Foundation (21-31712S) and the Basler Stiftung fuer Experimentelle Zoologie, and FC by an Australian Research Council (ARC) DECRA Fellowship (DE200100620).

## References

Andrews S. 2017. FastQC: a quality control tool for high throughput sequence data. 2010.

Baylor DA. 1987. Photoreceptor signals and vision. Proctor lecture. Investigative ophthalmology & visual science 28.1

Betancur-R R., et al. 2017. Phylogenetic classification of bony fishes. BMC evolutionary biology, 17(1), 162.

Biagioni LM, Hunt DM, Collin SP. 2016. Morphological characterization and topographic analysis of multiple photoreceptor types in the retinae of mesopelagic hatchetfishes with tubular eyes. Frontiers in Ecology and Evolution, 4, 25.

Britt LL, Loew ER, McFarland N. 2001. Visual pigments in the early life stages of Pacific northwest marine fishes. Journal of Experimental Biology 204.14.

Bowmaker JK, Hunt DM, Jeffery G. 2008. Eel visual pigments revisited: The fate of retinal cones during metamorphosis. Visual neuroscience, 25(3), 249.

Bozzano A, Pankhurst PM, Sabatés A. 2007. Early development of eye and retina in lanternfish larvae. Visual neuroscience, 24(3), 423–436.

Byun JH, et al. 2020. Gene expression patterns of novel visual and non-visual opsin families in immature and mature Japanese eel males. PeerJ, 8, e8326.

Carleton KL, Kocher TD. 2001. Cone opsin genes of African cichlid fishes: tuning spectral sensitivity by differential gene expression. Molecular biology and evolution 18.8.

Carleton KL, Dalton BE, Escobar‐Camacho D, Nandamuri SP. 2016. Proximate and ultimate causes of variable visual sensitivities: insights from cichlid fish radiations. Genesis, 54(6), 299–325.

Carleton KL, Escobar-Camacho D, Stieb SM, Cortesi F, Marshall NJ. 2020. Seeing the rainbow: mechanisms underlying spectral sensitivity in teleost fishes. Journal of Experimental Biology, 223(8).

Collin SP, Hoskins RV, Partridge JC. 1998. Seven Retinal Specializations in the Tubular Eye of the Scopelarchus michaelsarsi: A Case Study in Visual Optimization. Brain, Behavior and Evolution, 1998(09), 291–314.

Collin SP, Marshall NJ. 2003. Sensory Processing in Aquatic Environments. Springer-Verlag New York.

Cortesi F, et al. 2015. Ancestral duplications and highly dynamic opsin gene evolution in percomorph fishes. Proceedings of the National Academy of Sciences 112.5.

Cortesi F, et al. 2016. From crypsis to mimicry: changes in colour and the configuration of the visual system during ontogenetic habitat transitions in a coral reef fish. Journal of Experimental Biology, 219(16), 2545–2558.

Cortesi F, et al. 2020. Visual system diversity in coral reef fishes. Seminars in Cell & Developmental Biology. Academic Press.

Cortesi F, et al. 2021. Multiple ancestral duplications of the red-sensitive opsin gene (LWS) in teleost fishes and convergent spectral shifts to green vision in gobies. bioRxiv: https://doi.org/10.1101/2021.05.08.443214.

Dalton BE, Lu J, Leips J, Cronin TW, Carleton KL. 2015. Variable light environments induce plastic spectral tuning by regional opsin coexpression in the African cichlid fish, *Metriaclima zebra*. Molecular ecology 24,16 (2015).

de Busserolles F, et al. 2017. Pushing the limits of photoreception in twilight conditions: The rod-like cone retina of the deep-sea pearlsides. Science advances, 3(11), eaao4709.

de Busserolles F, Fogg L, Cortesi F, Marshall J. 2020. The exceptional diversity of visual adaptations in deep-sea teleost fishes. Seminars in Cell & Developmental Biology. Academic Press.

Dalton BE, Loew ER, Cronin TW, Carleton KL. 2014. Spectral tuning by opsin coexpression in retinal regions that view different parts of the visual field. Proceedings of the Royal Society B: Biological Sciences 281,1797.

Denton EJ. 1990. Light and vision at depths greater than 200 metres. Light and life in the sea 127–148.

Denton EJ, Herring PJ, Widder EA, Latz MF, Case JF. 1985 The roles of filters in the photophores of oceanic animals and their relation to vision in the oceanic environment. Proceedings of the Royal Society of London. Series B. Biological Sciences 225, 1238.

Douglas RH, Hunt DM, Bowmaker JK. 2003. Spectral sensitivity tuning in the deep-sea. In Sensory Processing in Aquatic Environments (pp. 323–342). Springer, New York, NY.

Douglas RH, Partridge JC, Marshall NJ. 1998. The visual systems of deep-sea fish. I. Optics, tapeta, visual and lenticular pigmentation. Prog. Ret. Eye Res, 17(4), 597–636.

Douglas RH, Genner MJ, Hudson AG, Partridge JC, Wagner HJ. 2016. Localisation and origin of the bacteriochlorophyll-derived photosensitizer in the retina of the deep-sea dragon fish *Malacosteus niger*. Scientific reports, 6, 39395.

Downes GB, Gautam N. 1999. The G protein subunit gene families. Genomics, 62(3), pp.544–552.

Enright JM, et al. 2015. *Cyp27c1* red-shifts the spectral sensitivity of photoreceptors by converting vitamin A1 into A2. Current Biology, 25(23), 3048–3057.

Hope AJ, Partridge JC, Dulai KS, Hunt DM. 1997. Mechanisms of wavelength tuning in the rod opsins of deep-sea fishes. Proceedings of the Royal Society of London. Series B: Biological Sciences, 264(1379), 155–163.

Huang Z, Titus T, Postlethwait JH, Meng F. 2019. Eye Degeneration and Loss of otx5b Expression in the Cavefish Sinocyclocheilus tileihornes. Journal of molecular evolution 87.7.

Hunt DM, Dulai KS, Partridge JC, Cottrill P, Bowmaker JK. 2001. The molecular basis for spectral tuning of rod visual pigments in deep-sea fish. Journal of Experimental Biology, 204(19), 3333–3344.

Hunt DM, Fitzgibbon J, Slobodyanyuk SJ, Bowmaker JK., Dulai KS. 1997. Molecular evolution of the cottoid fish endemic to Lake Baikal deduced from nuclear DNA evidence. Molecular Phylogenetics and Evolution, 8(3), 415–422.

Hunt DM, Hankins MW, Collin SP, Marshall NJ. 2014. Evolution of visual and non-visual pigments (Vol. 4). Boston, MA: Springer.

Katoh K, Asimenos G, Toh H. 2009. Multiple alignment of DNA sequences with MAFFT. In Bioinformatics for DNA sequence analysis (pp. 39–64). Humana Press.

Kawamura S, Tachibanaki S. 2012. Explaining the functional differences of rods versus cones. Wiley Interdisciplinary Reviews: Membrane Transport and Signaling 1.5.

Kearse M, et al. 2012. Geneious Basic: an integrated and extendable desktop software platform for the organization and analysis of sequence data. Bioinformatics, 28(12), 1647–1649.

La Vail MM, Rapaport DH, Rakic P. 1991. Cytogenesis in the monkey retina. J. Comp. Neurol. 309, 86–114. doi:10.1002/cne.903090107.

Lamb TD. 2013. Evolution of phototransduction, vertebrate photoreceptors and retina. Progress in retinal and eye research, 36, 52–119.

Lamb TD. 2019. Evolution of the genes mediating phototransduction in rod and cone photoreceptors. Progress in Retinal and Eye Research, p.100823.

Larhammar D, Nordström K, Larsson TA 2009.. Evolution of vertebrate rod and cone phototransduction genes. Philosophical Transactions of the Royal Society B: Biological Sciences, 364(1531), pp.2867–2880.

Liu D, et al. 2019. The cone opsin repertoire of osteoglossomorph fishes: gene loss in mormyrid electric fish and a long wavelength-sensitive cone opsin that survived 3R. Molecular biology and evolution 36.3.

Lythgoe JN. 1966. Visual pigments and under- water vision. In: Light as an Ecological Factor (Rackham, O., ed.). Oxford: Blackwell.

Luehrmann M, et al. 2019. Cardinalfishes (Apogonidae) show visual system adaptations typical of nocturnally and diurnally active fish. Molecular ecology 28.12.

Luo DG, et al. 2020. Apo-Opsin and Its Dark Constitutive Activity across Retinal Cone Subtypes.” Current Biology 30.24.

Ma JX, et al. 2001. A visual pigment expressed in both rod and cone photoreceptors. Neuron, 32(3), 451–461.

Mariani AP. 1986. Photoreceptors of the larval tiger salamander retina. Proceedings of the Royal society of London. Series B. Biological sciences 227.1249.

Mears AJ, et al. 2001. *Nrl* is required for rod photoreceptor development. Nature genetics, 29(4), 447–452.

Moser HG, Smith PE. 1993. Larval fish assemblages and oceanic boundaries. Bull. Mar. Sci, 53(2), 283–289.

Munk O. 1990. Changes in the visual cell layer of the duplex retina during growth of the eye of a deep‐sea teleost, Gempylus serpens Cuvier, 1829. Acta Zoologica, 71(2), 89–95.

Musilova Z, et al. 2019a. Vision using multiple distinct rod opsins in deep-sea fishes. Science, 364(6440), 588–592.

Musilova Z, et al. 2019b. Evolution of the visual sensory system in cichlid fishes from crater lake Barombi Mbo in Cameroon. Molecular Ecology, (August), 5010–5031.

Musilova Z, Salzburger W, Cortesi F. 2021. The Visual Opsin Gene Repertoires of Teleost Fishes: Evolution, Ecology and Function. Annual Review of Cell and Developmental Biology, 37, https://doi.org/10.1146/annurev-cellbio-120219-024915.

Musilova Z, Cortesi F. 2021. Multiple ancestral and a plethora of recent gene duplications during the evolution of the green sensitive opsin genes (RH2) in teleost fishes. bioRxiv, https://doi.org/10.1101/2021.05.11.443711.

Partridge JC, Shand J, Archer SN, Lythgoe JN, van Groningen-Luyben WA. 1989. Interspecific variation in the visual pigments of deep-sea fishes. Journal of Comparative Physiology A, 164(4), 513–529.

Partridge JC, Archer SN, van Oostrum J. 1992. Single and multiple visual pigments in deep-sea fishes. J. Mar. Biol. Assoc. U. K. 72, 113–130

Pointer MA, Carvalho LS, Cowing JA, Bowmaker JK, Hunt DM. 2007. The visual pigments of a deep-sea teleost, the pearl eye Scopelarchus analis. Journal of Experimental Biology, 210(16), 2829–2835.

Raymond PA. 1995. Development and morphological organization of photoreceptors. In Neurobiology and Clinical Aspects of the Outer Retina (pp. 1–23). Springer, Dordrecht.

Reif WE. 1985. Functions of scales and photophores in mesopelagic luminescent sharks. Acta Zoologica, 66(2), 111–118.

Ronquist F, et al. 2012. MrBayes 3.2: efficient Bayesian phylogenetic inference and model choice across a large model space. Systematic biology, 61(3), 539–542.

Sassa C, Hirota Y. 2013. Seasonal occurrence of mesopelagic fish larvae on the onshore side of the Kuroshio off southern Japan. Deep Sea Research Part I: Oceanographic Research Papers, 81, 49–61.

Savelli I, Flamarique IN, Iwanicki T, Taylor JS. 2018. Parallel opsin switches in multiple cone types of the starry flounder retina: tuning visual pigment composition for a demersal life style. Scientific reports 8:1.

Schott RK, Bhattacharyya N, Chang BS. 2019. Evolutionary signatures of photoreceptor transmutation in geckos reveal potential adaptation and convergence with snakes. Evolution, 73(9), 1958–1971.

Schott RK, et al. 2016. Evolutionary transformation of rod photoreceptors in the all-cone retina of a diurnal garter snake. Proceedings of the National Academy of Sciences, 113(2), 356–361.

Sernagor E, Eglen S, Harris B, Wong R. 2006. Retinal development. Cambridge University Press.

Shen YC, Raymond PA. 2004. Zebrafish cone-rod (*crx*) homeobox gene promotes retinogenesis. Developmental biology, 269(1), 237–251.

Simoes BF, et al. 2016. Multiple rod–cone and cone–rod photoreceptor transmutations in snakes: evidence from visual opsin gene expression. Proceedings of the Royal Society B: Biological Sciences, 283(1823), 20152624.

Stieb SM, et al. 2019. A detailed investigation of the visual system and visual ecology of the Barrier Reef anemonefish, *Amphiprion akindynos*. Scientific reports 9.1.

Sugawara T, et al. 2005. Parallelism of amino acid changes at the RH1 affecting spectral sensitivity among deep-water cichlids from Lakes Tanganyika and Malawi. Proceedings of the National Academy of Sciences, 102(15), 5448–5453.

Tettamanti V, de Busserolles F, Lecchini D, Marshall NJ, Cortesi F. 2019. Visual system development of the spotted unicornfish, *Naso brevirostris* (Acanthuridae). Journal of Experimental Biology, 222(24).

Torres-Dowdall J, et al. 2017. Rapid and parallel adaptive evolution of the visual system of Neotropical Midas cichlid fishes. Molecular biology and evolution 34.10.

Turner JR, White EM, Collins MA, Partridge JC, Douglas RH. 2009. Vision in lanternfish (Myctophidae): adaptations for viewing bioluminescence in the deep-sea. Deep Sea Research Part I: Oceanographic Research Papers 56, 6.

Underwood G. 1968. Some suggestions concerning vertebrate visual cells. Vision research, 8(4), 483–488.

Valen R, et al. 2016. The two-step development of a duplex retina involves distinct events of cone and rod neurogenesis and differentiation. Developmental biology, 416(2), 389–401.

Wagner HJ, Partridge JC, Douglas RH. 2019. Observations on the retina and ‘optical fold’of a mesopelagic sabretooth fish, *Evermanella balbo*. Cell and Tissue Research, 378(3), 411–425.

Yokoyama S. 2008. Evolution of dim-light and color vision pigments. Annu. Rev. Genomics Hum. Genet., 9, 259–282.

Yokoyama S, Jia H. 2020. Origin and adaptation of green‐sensitive (RH2) pigments in vertebrates. FEBS Open Bio, 10(5), 873–882.

Zhang H, et al. 2020. Molecular cloning of fresh water and deep‐sea rod opsin genes from Japanese eel *Anguilla japonica* and expressional analyses during sexual maturation. Febs Letters, 469(1), 39–43.

