## Supplementary material for "Visual gene expression reveals a cone to rod developmental progression in deep-sea fishes": Supp file 1

**Supplementary file 1: Genomic sequences**

Chauliodus_sloani_56H1_RH2_genome_var01_FIN: ATGAACGGCACAGAGGGAAGCAACTTCTACATCCCCATGTCCAACCGGACTGGGCTTGTCAGGAGTCCGTATCACTACCCACAGTACTACCTGGGAGCTCCATATCAATTCTACATGCTTGCGGCCTACATGTTCTTCCTGATTTGTTTCGGTTTCCCCATCAACGCTCTGACACTGGTGGTCACAGCTCAGAACAAGAAGCTCCGGCAGCCCCTCAATTTCATCCTGGTCAACTTGGCTGTGGCTGGACTCATCATGGTCTGCTTTGGCTTCACCATGACCTTCATCACTGCTATTAATGGCTACTTCATCTTTGGGCCAATGGGTTGTGCCATTGAGGGCTTCATGGCTACTCTTGGAGGTCAGGTTTCCCTGTGGTCCTTGGTTGTGCTGGCTATCGAGAGATACATTGTTGTCTGCAAACCCATGGGCAGCTTCAAGTTTGGAGCCGCCCATGCAGGTGTTGGAGTGGTTTTCACCTGGATCATGGCCATGGCCTGCGCTGCTCCCCCCCTGGTTGGCTGGTCCAGGTACCTACCCGAGGGCCTGCAGTGCTCCTGTGGACCAGACTACTACACCCTCAACCCTGAATTCAACAATGAATCCTACGTCATGTACATGTTCTCCTGCCACTTCTGTGTCCCTGTGTTCACCATCTTCTTCACTTATGGAAGCCTTGTGTTGACAGTCAAAGCGGCTGCCGCTTCCCAGCAGGAGTCAGAGTCCACTCAGAAGGCTGAGAGGGAAGTGACACGTATGTGCTTCCTGATGGTCATGGGCTTCCTGATTGCATGGGTTCCCTATGCCAGCTTTGCTGCTTACATCTTCATGAACAAAGGCGTAGCCTTCACTGCCCAGGCCATGACTGTCCCTGCCTTCTTCTCCAAGGCCTCAGCTCTGTTCAACCCTATTATCTATGTGCTCATGAACAAACAGTTCCGTAGCTGCATGCTGTCAACTTTGGGTATGGGCGGTATGGTTGATGATGAGAGCTCAGTTTCTTCATCAAGCAAGACAGAAGTGTCTTCTGTGTCTACAGCATAA

Chauliodus_sloani_56H1_RH2_genome_var02_FIN: ATGAACGGCACAGAGGGAAGCAACTTCTACATCCCCATGTCCAACCGGACTGGGCTTGTCAGGAGTCCGTATCACTACCCACAGTACTACCTGGGACCTCCATATCAATTCTACATGCTTGCGGCCTACATGTTCTTCCTGATTTGTTTCGGTTTCCCCATCAACGCTCTGACACTGGTGGTCACAGCTCAGAACAAGAAGCTCCGGCAGCCCCTCAATTTCATCCTGGTCAACTTGGCTGTGGCTGGACTCATCATGGTCTGCTTTGGCTTCACCATGACCTTCATCACTGCTATTAATGGCTACTTCATCTTTGGGCCAATGGGTTGTGCCATTGAGGGCTTCATGGCTACTGTTGGAGGTCAGGTTTCCCTGTGGTCCTTGGTTGTGCTGGCTATCGAGAGATACATTGTTGTCTGCAAACCCATGGGCAGCTTCAAGTTTGGAGCCGCCCATGCAGGTGCTGGAGTGGCTTTCACCTGGATCATGGCCATGGCCTGCGCTGCTCCCCCGCTGGGTGGCTGGTCCAGGTACATACCCGAGGGCCTGCAGTGCTCCTGTGGACCAGACTACTACACCCTCAACCCTGAATATAACAACGAATCCTACGTCATGTACATGTTCTCCTGCCACTTCTGTGTCCCTGTGTTCACCATCTTCTTCACTTATGGAAGCCTTGTGTTGACAGTCAAAGCGGCTGCCGCTTCCCAGCAGGAGTCAGAGTCCACTCAGAAGGCTGAGAGGGAAGTGACACGTATGTGCGTCCTGATGGTCTTGGGCTTCCTGGTTGCATGGGTTCCCTATGCCAGCTTTGCTGCTTACATCTTCCTGAACAAAGGCGTAGCCTTCTCTGCCCAGGCCATGACTGTCCCTGCCTTCTTCTCCAAGGCCTCAGCTCTGTTCAACCCTATTATCTATGTGCTCATGAACAAACAGTTCCGTAACTGCATGCTGACAACTGTGGGTATGGGCGGTATGGTTGAAGATGAGAGCTCAGTTTCTTCATCAAGCAAGACAGAAGTGTCTTCAGCATAA

Coccorella_atlantica_56C7_RH2_genome_var01_adult_FIN: ATGGAGAACGGCACAGAGGGCAAGAACTTCTACGTTCCTATGAACAATAGGAGTGGGCTTGTGAGGAGTCCTTTTGAATACCCACAGTATTACTTGGCAGATCCAATGCTGTACAAGTTTCAGGCTTTCTACATGCTTTTCCTCATCTTTACCGGCGGCCCCATCAATTTCCTGACACTGTTTGTCACGGTTCAGAACAAGAAACTCCGGCAACCACTTAACTTCATTCTGGTCAACTTGGCGGTCGCYGGCGCAATCATGGTGGTTTTTGGATTCACAGTCACCTTTTATACTTCCATGTGTGGCTACTTTGCTTTGGGACCAGCCAGCTGTGCTGCGGAGGGTTTCTTTGCTACAATTGGAGGTCAGGTAGCTCTGTGGTCTCTTGTGGTTCTTGCCGTTGAGAGGTACATTGTGGTCTGCAAACCCATGGGYAGCTTCAAATTCACKGCCACGCACGCCGGCGCCGGATGTGCACTCACATGGGTCATGGCCATGTCTTGTGCTGCACCACCTCTCGTTGGCTGGTCCAGGTACCTCCCCGAGGGCTTGCAGTGCTCCTGTGGACCCGACTACTACACCCTGGCCCCGGGCTTCAACAACGAGTCCTACGTCTTGTACATGTTCAGCTGTCACTTCACGGTGCCCCTCGTCACCATCTTCTTCACTTATGGAAGCCTCGTGCTGACAGTCAAAGCGGCCGCAGCTCAGCAGCAGGAGTCAGAGTCCACCCAGAAGGCCGAGAGGGAAGTGACACGTATGTGTATCGTGATGGTCATTGGCTTCCTGGTGACATGGGTTCCATACGCCAGTCTCGCTGCTTGGATCTTCTTCAACCGGGGAGCTGCCTTCTCTGCTACTGGCATGGCTGTTCCTGCTTTCTTCGCCAAAGCCTCAGCCTTGTTCAACCCAATCATCTATGTGTTGTTGAACAAACAGTTCCGTAACTGCATGCTGAGCACCGTTGGAATGGGTGGCATGGTAGAAGATGAGGGAGCTTCCTCTACCAGCAGCAAGACAGAAGTCTCCCAAGGATAA

Coccorella_atlantica_56C7_RH2_genome_var02_larval_FIN: ATGGAGAACGGCACAGAGGGCAAGAACTTCTACATCCCCATGAACAACAGGACGGGGCTTGCGAGGAGTCCTTTTGAGTACCCACAGTATTACCTGGCGGATCCCATGGTGTTCAAGTTACAGGCTGCTTACATGTTTTTCCTCATCATTCTTGGTGGTCCCCTGAACGGCCTGACGCTGGTGGTCACGGCTCAGAACAAGAAGCTCCGGCAACCTCTTAACTTTATTCTGGTCAACTTGGCGGTCGCTGGATTGATCATGGTCATCGGTGGATTCACAGTCACATTCGTAACCGCAATGAATGGCTACTTTATCTTTGGACCGATGGGCTGTGCCTTGGAGGGAGGACTGGCTACGCTAGGAGGTCAGATCTCTCTGTGGTCCTTGGTAGTGCTGGCTGTTGAGAGGTACATTGTTGTCTGCAAACCCATGGGCAGCTTCAAGTTCACTGCAACTCACGCTGGAGTTGGCGTGGTGTTCACCTGGATCATGGCCGCTGCTTGCGCTTTCCCCCCATTCTTCGGCTGGTCCAGGTATATCCCAGAGGGAATCCAGACCTCATGTGGACCTGACTACTACACCATGGCCCCAGGCTTCAACAACGAATCATACGTCATGTACATGTTCAGCTGTCACTTCTGTTTCCCAGTCTTCACCATCTTCTTCACTTATGGAAGCCTGGTGCTGACAGTCAAAGCGGCTGCAGCTCAGCAGCAGGAGTCAGAATCCACCCAGAAAGCTGAGAGGGAAGTCACACGTATGTGCGTCCTGATGGTCTTGGGCTTCCTGACTGCCTGGGTTCCATATGCCAGCTATGCCGCCTGGATCTTCTTCAACAAGGGAGCTGCCTTCTCTGCTATCTCCATGGCCATCCCTGCCTTCTTTTCCAAGGCCTCAGCCGTGTTCAACCCAATCATCTATGTGCTGTTGAACAAACAGTTCCGTAACTGCATGCTCAGCACTATTGGAATGGGAGGCATGGTTGATGATGAGAGCTCAGTGTCTGGCAGCAGCAAGACAGAAGTCTCCTCAGTTTCTCAAGGCTAG

Coccorella_atlantica_56C7_RH2_genome_var03_FIN: ATGGAGAACGGCACAGAGGGCAAGAACTTCTACATCCCCATGAACAACAGGACGGGGCTTGCGAGGAGTCCTTTTGAGTACCCACAGTTTTACCTGGCGGATCCCATGGTGTTCAAGTTTCAGGCTTTCTACATGTTCATCCTCATCATGCTCGGYGGCCCCATCAACGGCCTGACGCTGCTGGTCACGGCTCAGAACAAGAAGCTCCGGCAACCTCTTAACTTTATTCTGGTCAACTTGGCGGTCGCTGGATTGATCATGGTCATGGGTGGATTCACCATCACATTCGTAACCGCAATCTATGGCTACTTTATTTTTGGACCGTTGGGCTGTGCCTTGGAGGGAGGACTGGCTACGCTAGGAGGTCAGGTTTCTCTGTGGTCCCTGGTAGTGCTGGCTGTTGAGAGATACATTGTTGTCTGCAAACCCATGGGCAGCTTCAAGTTCACTACGACTCACGCTGCAGTTGGAGTTGCTTTCACCTGGATCATGGCTTCGTCTTGTGCTTTGCCTCCATTCTTCGGCTGGTCCAGGGTATATTCCAGAGGGAATCCAGACCTCATGTGGACCTGACTACTATACCCAGGCCCCAGGCTACAACAACGAATCATATGTCTTGTACCTGTTCAGCTGTCACTTCTGTGTCCCTGTCGTCACCATCTTCTTCACTTATGGAAGCCTTGTGCTGACAGTCAAAGCGGCTGCAGCTCAGCAGCAGGAGTCAGAATCCACCCAGAAAGCTGAGAGGGAAGTCACACGTATGTGCATCCTGATGGTCTGTGGCTTCCTGCTTGCCTGGACTCCATATGCCAGCTTTGCAGCGTGGATCTTCTTCAACAAGGGAGCTGCCTTCACTGCTACCGGCATGGCCATCCCTGCCTTCTTCTCCAAGTCCTCAGCCTTGTTTAACCCAATCATCTATGTGCTGATGAACAAACAGTTCCGTAACTGCATGCTGACCACTGTAGGAATGGGCGGCATGGTAGAAGATGAGACCTCAGTGTCCACCAGCAAGACTGAAGTCTCCTCTGTGTCTTAA

Scopelarch_71B7_RH2_adult_RH2_ex1_var01a_corrected_X: ATGGAGAACGGCACAGAGGGCAAGAACTTCTATATCCCCATGAACAACAGGAGTGGGCTCGTGAGGAGTCCCTATGAATACCCACAGTACTACCTGGCCGATCCCATAATATTCAAGTTACAGGCTGCCTATATCCTTTTCCTGATGTTCACCGGCGGACCCATCAATGTTCTAACCTTGGTGGTCACAGCTCGGAACAAGAAGCTCCAGCAACCTCTTAACTTYATTCTGGTCAACTTGGCTTTCGCTGGAGCACTTATGGTGTTTGGTGGATTTGTAATGACCTTTTATACTTCAATGAATGGCTACTTTGTTTTGGGACCRATGAGCTGTGCTTTTGAGGGCTTCATGGCCACACTTGGAGCCTGTGGTCCTGTGCGGTTCTGGCCGTTGAGAGATACATTGTGGTCTGCAAACCTTTGGGTAACTTCAAATTCTCGGCCTCTCACGCCGCCA

Scopelarch_71B7_RH2_adult_RH2_ex1_var01b_VI: ATGGAGAACGGCACAGAGGGCAAGAACTTCTATATCCCCATGAACAACAGGAGTGGGCTCGTGAGGAGTCCCTATGAATACCCACAGTACTACCTGGCCGATCCCGTAATGTTCAAGATCCAGGCTTTCTACATGTTTTTCGTGATTGTCACCGCCGGACCCATCAATATTCTGACCATGGTGGTCACAGCTCAGAACAAGAAGCTCCGGCAACCAATTAACTACATGCTGGTCAACTTGGCTTTCGCTGGAGCAATTATGGTGTTTGGTAATTTAATCTGCTTTTATTCTTCAATGCATGGCTACTTTCCTTTGGGACCGATCAGCTGTGCTATTGAGGGCTTCACGGCCACAATTGGAGCCTGTGGTCCCTCGTGGTTCTGGCCGTTGAGAGATACATTGTGGTCTGCAAACCTTTGGGTAACTTCAAATTCTCGGCCTCTCACGCCGCCA

Scopelarch_71B7_RH2_adult_RH2_ex1_var01c_VI: ATGGAGAACGGCACAGAGGGCAAGGACTTCTATATCCCCATGAACAACAGGAGTGGGCTCGTGAGGAGTCCCTATGAATACCCACAGTACTACCTGGCCGATCCCATAATGTTCAAGTTCCAGGCTTTCTACATGCTTTTCCTGATGCTCACCGGCGGACCCATCAATATTCTGACCTTGGTGGTCACAGCTCAGAACAAGAAGCTCCGGCAACCTCTTAACTTCATTCTGGTCAACTTGGCTGTCGCTGGAGCAATTATGGTGTTTGGTGGATTTTTAATCACCTTTTATACTTCAATGAATGGCTACTTTCTTTTGGGACCGATGAGCTGTGCTATTGAGGGCTTCATGGCCACACTTGGACCCTGTGGTCCCTCGTGGTTCTGGCCGTTGAGAGATACATTGTGGTCTGCAAACCCATGGGTAGCTTCAAATTCTCGGCCACTCACGCCGGGA

Scopelarch_71B7_RH2_adult_RH2_ex1_var01d1_VI: ATGGAGAACGGCACAGAGGGCAAGAACTTCTATATCCCCATGAACAACAGGAGTGGGCTCGTGAGGAGTCCCTATGAATACCCACAGTACTACCTGGCCGATCCCATATTCTTCAAGTTACAGGCTGCCTACATCCTTTTCCTGATGTCCACCGGCGGACCCATCAATATTCTGACCTTGGTGGTCACAGCTCAAAACAAGAAGCTCCAGCAACCTCTTAACTTCATTCTGGTCAACTTGGCTTTCGCTGGAGCACTTATGGTGTTTGGTGGATTTGTAATGACCTTTTATACTTCAATGAATGGCTACTTTGTTTTGGGACCGATGAGCTGTGCTTTTGAGGGCTTCATGGCCACACTTGGACCCTGTGGTCCCTYGTGGTTCTGGCCGTTGAGAGATACATTGTGGTCTGCAAACCCATGGGTAGCTTCAAATTCTCGTCCACTCACGCCGGGA

Scopelarch_71B7_RH2_adult_RH2_ex1_var01d2_VI: ATGGAGAACGGCACAGAGGGCAAGAACTTCTATATCCCCATGAACAACAGGAGTGGGCTCGTGAGGAGTCCCTATGAATACCCACAGTACTACCTGGCCGATCCCATAATATTCAAGTTACAGGCTGCCTACATCCTTTTCCTGATGTCCACCGGCGGACCCATCAATATTCTGACCTTGGTGGTCACAGCTCAAAACAAGAAGCTCCAGCAACCTCTTAACTTCATTCTGGTCAACTTGGCTTTCGCTGGTGCACTTATGGTGTTTGGTGGATTTGTAATGACCTTTTATACTTCAATGAATGGCTACTTTGTTTTGGGACCGATGAGCTGTGCTTTTGAGGGCTTCATGGCCACACTTGGACCCTGTGGTCCCTYGTGGTTCTGGCCGTTGAGAGATACATTGTGGTCTGCAAACCCATGGGTAGCTTCAAATTCTCGTCCACTCACGCCGGGA

Scopelarchus_RH2_adult_ex1complete_var02_VI: ATGGAGAACGGCACAGAGGGCAAGGACTTCTATATCCCCATGAACAACAGGAGTGGGCTCGTGAGGAGTCCCTATGAATACCCACAGTACTACCTGGCCGATCCCGTAATGTTCAAGTTCCAGGCTCTCTTCATGTTTTTCGTGATTATCACCGCCGGACCCATCAATATTCTGACCTTGGTGGTCACAGCTCAGAACAGGATGCTCCGGCAACCTATTAACTACATTATGCTCAACTTGGCTGTCGCTGGAATAATTATGGTGTTTAGTGGAAATTTTATCGCCTTTTATTGTTCAATGCATGGCTACTTTATTTTGGGACCGATGGGCTGTGCTATTGAGGGCTTCTCGGCCACAATTGGACCCTGTGGTCCCTCGTGGTTCTGGCCGTTGAGAGATACATTGTGGTCTGCAAACCCATGGGTAGCTTCAAATTCTCGTCCACTCACGCCGGGA

Scopelarchus_RH2_genome_var03_FIN: ATGGAGAACGGCACAGAGGGCAAGAACTTCTACATTCCCATGAACAACAGGACTGGGCTTGTGAGGAATCCTTTCGAATACCCACAGTATTACTTGGCAGATCCAATGGTGTTCAAGTTACAGGCAGCCTACATGTTTTTCCTCATCATGTTTGGCGGCCCCATCAACGGTCTGACATTGGTGGTCACAGCTCAGAACAAGAAGCTCCGGCAACCTCTTAACTTCATTCTGGTCAACTTGGCTGTCGCTGGAATGATCATGGTCCTTGGTGGATTCACCATCACCTTCATAACTGCTATTAATGGCTACTTTGTTTTTGGACCATTCGCATGTGCCCTTGAGGGAGGGCTGGCTACACTAGGAGGTCAGATTTCTCTGTGGTCACTGGTAGTGCTGGCTGTTGAGAGATACATTGTCGTCTGTAAACCCATGGGAAGCTTCAAGTTCACTACAACTCATGCTGCCATTGGAGTTGTTTTCACCTGGATCATGGCTTCTGCTTGTGCTTTCCCTCCCTTCTTTGGCTGGTCCAGGTATATTCCCGAGGGAATCCAGACCTCATGTGGACCTGACTACTACACCATGGCCCCTGGCTTCAACAATGAATCATATGTCATGTACATGTTCAGCTGCCACTTCTGCTTCCCTGTCTTCACCATCTTCTTCACTTATGGAAGCCTTGTCCTGACAGTCAAAGCCGCTGCAGCTCAGCAGCAGGAGTCAGAATCCACCCAGAAAGCTGAGAGGGAAGTCACACGTATGTGCGTCTTGATGGTCTTTGGCTTCCTGACTGCATGGGTACCATATGCCAGCTATGCTGCTTATATCTTCTTCAACAAGGGAGTAGCCTTCTCTGCCGTGTCCATGGCCATCCCTGCCTTCTTCTCCAAGGCCTCAGCTGTGTTCAACCCAGTCATCTATGTGCTCATGAACAAACAGTTCCGTAACTGCATGCTCAGCACTATTGGAATGGGTGGCATGGTTGATGATGAGAGTTCAGTGAGTGGCTCCTCCAGCAGCAAGACAGAAGTCTCCTCTGTTTCTCAAGGATAG

Scopelarchus_RH2_genome_var05_FIN: ATGGAGAACGGCACAGAGGGCAAGAACTTCTACATCCCCATGAACAACAGGACTGGGCTTGTGAGGAGTCCTTTCGAATACCCACAGTATTACTTGGCAGATTCAATGGTGTTCAAGTTTCAGGCAGCCTACATGTTTTTCCTCATCATGTTTGGCGGCCCCATCAACGGTCTGACATTGGTGGTCACAGCTCAGAACAAGAAGCTCCGGCAACCTCTTAACTTCATTCTGGTCAACTTGGCTGTCGCTGGAATGATCATGGTCCTTGGTGGATTCACCGTCACCTTCGTAACTGCTATTAATGGCTACTTTGTTTTTGGACCATTGGGATGTACCCTTGAGGGAGGGCTGGCTACATTAGGAGGTCAGGTTTCTCTCTGGTCACTGGTAGTGCTGGCTGTTGAGAGATACATTGTTGTCTGCAAACCTATGGGCAGCTTCAAGTTCACTACAACTCATGCTGCCATTGGAGTTGTTTTCACCTGGATCATGGCTTCTTCTTGTGCTCTCCCTCCCTTCTTTGGCTGGTCCAGGTATATTCCAGAGGGAATCCAGACCTCATGTGGACCTGACTACTACACCATGGCCCCTGGCTACAACAATGAATCATTTGTCATGTACATGTTCAGCTGCCACTTCTGTTTCCCCGTCTTCACCATCTTCTTCACCTATGGAAGCCTTGTCCTGACAGTCAAAGCGGCTGCAGCTCAGCAGCAGGAGTCAGAATCCACCCAGAAAGCTGAGAGGGAAGTCACACGTATGTGYGTCTTGATGGTCTTTGGCTTCCTGACTGCCTGGACACCATATGCCAGCTATGCTGCTTGGATCTTCTTCAACAAGGGAGCTGCCTTCTCTGCCGTGTCCATGGCCATCCCTGCCTTCTTCTCCAAGGCCTCAGCTGTGTTCAACCCAGTCATCTATGTGCTCATGAACAAACAGTTCCGTAACTGCATGCTGACCACTATTGGAATGGGTGGCATGGTTGATGATGAGACCTCAGTTTCCAGCAGCAGCAAGACAGAAGTCTCCTCAGTTTCTCAAGGATAG

Scopelarchus_RH2_var01_ex345_X: TACATTCCCGAGGGCCTGCAGTGCTCCTGTGGCCCTGACTACTACACCATGGCCCCTGGCTTCAACAATGAATCATATGTCATGTACATGTTCAGCTGCCACTTCTGCTTCCCCSTCTTCACCATCTTCTTCACTTATGGAAGCCTTGTCCTGACAGTCAAAGCGGCCGCAGCTCAGCAGCAGGAGTCAGAATCCACACAGAAGGCTGAGAGGGAAGTCACGCGTATGTGCTGTGTGATGGTCTTGGGCTTCCTCACTGCCTGGGTACCATACGCCAGTCTTGCTGCTTGGATCTTCTTCAACAAGGGAGCTGCCTTCTCTGCCGTTGGCATGGCTGTTCCTGCTTTCTTCGCAAAAGCCGCAGCCTTGTTCAACCCAATCATCTACATCTTGTTCAACAAACAGTTCCGTAACTGCATGCTGACTACTATTGGAATGGGTGGCATGGTTGATGATGAGAGCTCAGTTTCCAGCAGCAGCAGCAAGACAGAAGTCTCCTCAGTTTCCCAAGGATAA

Scopelarchus_RH2_var02_ex345_X: TACGTTCCCGAGGGCCTGCAGTGCACCTGTGCATTTGACTACTACACCCTGGCCCCTGGCTTCAACAATGAATCATTTGTCATGTTCATCTTCTGCTGCCACTTCTGCTTCCCCCTCTTCACCATCTTCTTCAGTTATGGAAGCCTTGTCCTGACAGTCAAATCGGCCGCAGCTCAGCAGGAGTCAGAATCCACACAGAAGGCTGAGAGGGAAGTCACGCGTATGTGCTGTGTGCTGGCCTTGGCCTGGCTCATTTTATGGCTACCATACGCCACTTTTTCTTTTTGGATCTTCTTCAACAAGGGAGCTGCCTTCTCTGCCGTTGGCAGGGCTGTTCCTACTTTCTACGCAAAACTCGCACCCTTGTTCCACATCATCATCTACATCTTGTTCAACAAACAGTTCCGTAACTCCATGCTGACTACTATTGGAATGAGTGGCATGGTTGATGATGAGAGCTCAGTGACCTCCAACAGCAGCAAGACAGATGTCTCCTCAGGATAA

Scopelarchus_RH2_var04a_ex345_X: TACATTCCCGAGGGCCTGCAATGCTCCTGTGGACCTGACTACTACACCCTGGCCCCTGGCTTCAACAATGAATCATATGTCATATACATGTTCACCTGCCACTTCATCTTCCCCGTCATCACCATCTTCTTCACTTATGGAAGCCTTGTCCTGACAGTCAAAGCGGCCGCAGCTCAGCAGCAGGAGTCAGAATCCACACAGAAGGCTGAGAGGGAAGTCACGCGTATGTGCTGTGTGATGGTCTTGGCCTTCCTCATTGCCTGGACCCCATACGCCAGTCTTACTGCTTGGATCTTCTTAAACAAGGGAGCTGCCTTCTCTGCTATTAGCATGGCTGTTCCTGCTTTCTTCGCAAAATCCTCATCCTTGTACAACCCAGTCATCTACGTGTTGTTGAACAAACAGTTCCGTAACTGCATGCTGTCTTCTATTGGAATGGGTGGCATGGTTGATGATGAGAGCTCAGTGACAAGCAGCAAGACTGAAGTCTCCTCAGTTTCCCAAGGATAA

Scopelarchus_RH2_var06_ex345_X: TACATTCCCGAGGGCCTGCAGTGCTCCTGTGGAATTGACTACTACACCATGGCCCCTGGCTTCAACAATGAATCATTTGTCATGTTCATCTTCTGCTGCCACTTCTGCTTCCCCCTCTTCACCATCTTCTTCACTTATGGAAGCCTTGTCCTGACAGTCAAAGCGGCCGCAGCTGAGCAGCAGGAGTCAGAATCCACACAGAAGGCTGAGAGGGAAGTCACGCGTATGTGCATTGTGATGGTCTTTGGCTGGCTCATTTTATGGATACCATACGCCAGTTTTGCTTTTTGGATCTTCTTCAACAAGGGAGCTGCCTTCTCTGCCGTTGGCAGGGCTGTTCCTTGTTTCTTCGCAAAACTCGCACCCTTGTTCCACATCCTCATCTACATCTTGTTCAACAAACAGTTCCGTAACTGCATGCTGGCTACTATTGGAATGGGTGGCATGGTTGATGATGAGAGCTCAGTGACCTCCACCAGCAGCAAGAAAGAAGTCTCCTCAGGATAA

Scopelarchus_RH2_var07_ex345_X: TACATTCCCGAGGGCCTGCAGTGCTCCTGTGGACCTGACTACTACACCCTGGCCCCTGGCTTCAACAATGAATCATATGTCATATACATGTTCACCTGCCACTTCATCTTCCCCGTCATCACCATCTTCTTCACTTATGGAAGCCTTGTCCTGACAGTCAAAGCGGCCGCAGCTCAGCAGCAGGAGTCAGAATCCACACAGAAGGCTGAGAGGGAAGTCACGCGTATGTGCTGTGTGATGGTCTTGGCCTTCCTCGTTGCCTGGACCCCATACGCCAGTCTTACTGCTTGGATCTTCTTAAACAAGGGAGCTGCCTTCTCTGCTATTAGCATGGCTGTTCCTGCTTTCTTCGCAAAATCCTCATCCTTGTACAACCCAATCATCTACGTGTTGTTGAACAAACAGTTCCGTAACTGCATGCTGTCTGCTATTGGAATGGGTGGCATGGTTGATGATGAGAGCTCAGTGACAAGCAGCAAGACTGAAGTCTCCTCAGTTTCCCAAGGATAA
